# Single-Cell Genome Dynamics in Early Embryo Development: A Statistical Thermodynamics Approach

**DOI:** 10.1101/123554

**Authors:** Alessandro Giuliani, Masa Tsuchiya, Kenichi Yoshikawa

## Abstract

A statistical thermodynamics approach to the temporal development of biological regulation provides a phenomenological description of the dynamical behavior of genome expression in terms of autonomous self-organization with a critical transition (Self-Organized Criticality: SOC). In early mouse embryo development, the dynamical change in the self-organization of overall expression determines *how and when* reprogramming of the genome-expression state occurs. Reprogramming occurs via a transition state (climbing over an epigenetic landscape), where the critical-regulation pattern of the zygote state disappears. A critical transition is well captured in terms of the bimodality of expression ensembles, which reflects distinct thermodynamic states (critical states). These critical states exhibit a genome avalanche pattern: competition between order (scaling) and disorder (divergence) around a critical point. The genome avalanche in mouse embryo development, which is committed to erase a previous ordered state, reveals that the direction of early embryo single-cell development traverses the same steps as in differentiation, but in the opposite order of self-organization.

## Introduction

A fundamental issue in bioscience is to understand the mechanism that underlies the dynamic control of genome-wide expression through the complex temporal-spatial self-organization of the whole genome to regulate changes autonomously in the cell fate.

Our recent studies on genome-scale expression dynamics in the cell-fate change demonstrated the possibility of a statistical thermodynamic analysis [1-3]. The thermodynamics phenomenological character is at the basis of Albert Einstein’s famous statement ‘*A theory is the more impressive the greater the simplicity of its premises is, the more different kinds of things it relates, and the more extended is its area of applicability. Therefore, the deep impression which classical thermodynamics made upon me. It is the only physical theory of universal content which I am convinced that, within the framework of applicability of its basic concepts, it will be never overthrown*’ [4].

An open-thermodynamic approach clearly reveals how and when the change in a genome-expression state occurs as a thermodynamic event in the disappearance of critical phenomena that arise from an initial state. Furthermore, a detailed global perturbation mechanism for the self-organization of overall expression was elucidated for cell differentiation processes. This demonstration of statistical thermodynamics in the genome [5] is particularly crucial in biology, where the extreme complexity of the system rules out any current strict mechanistic approach that considers all of the microscopic regulations involved.

The active process that controls global gene expression in the case of cell fate determination, involves a particular kind of self-organization, called Self-Organized Criticality (SOC) [2, 6-8], which drives the state transitions of gene expression via the mutual interaction among three different gene expression states; ‘critical’, ‘near critical’ and ‘sub-critical’. Bimodality in gene expression levels is a clear signature of the self-organizing critical transition that is organized into distinct critical states (**Figure 1**). Each gene belongs to one of these three states based on the relative variation of its expression value. This allows for a phenomenological (and thus independent of any mechanistic hypothesis) description of the expression dynamics at the global level.

In this report, we investigate whether essentially the same critical-state dynamics we observed for cell differentiation processes [2,3] does exist in overall RNA expression of single-cell mouse embryo development. This case is particularly relevant to give a further proof of SOC control as universal character.

**Figure 1:**
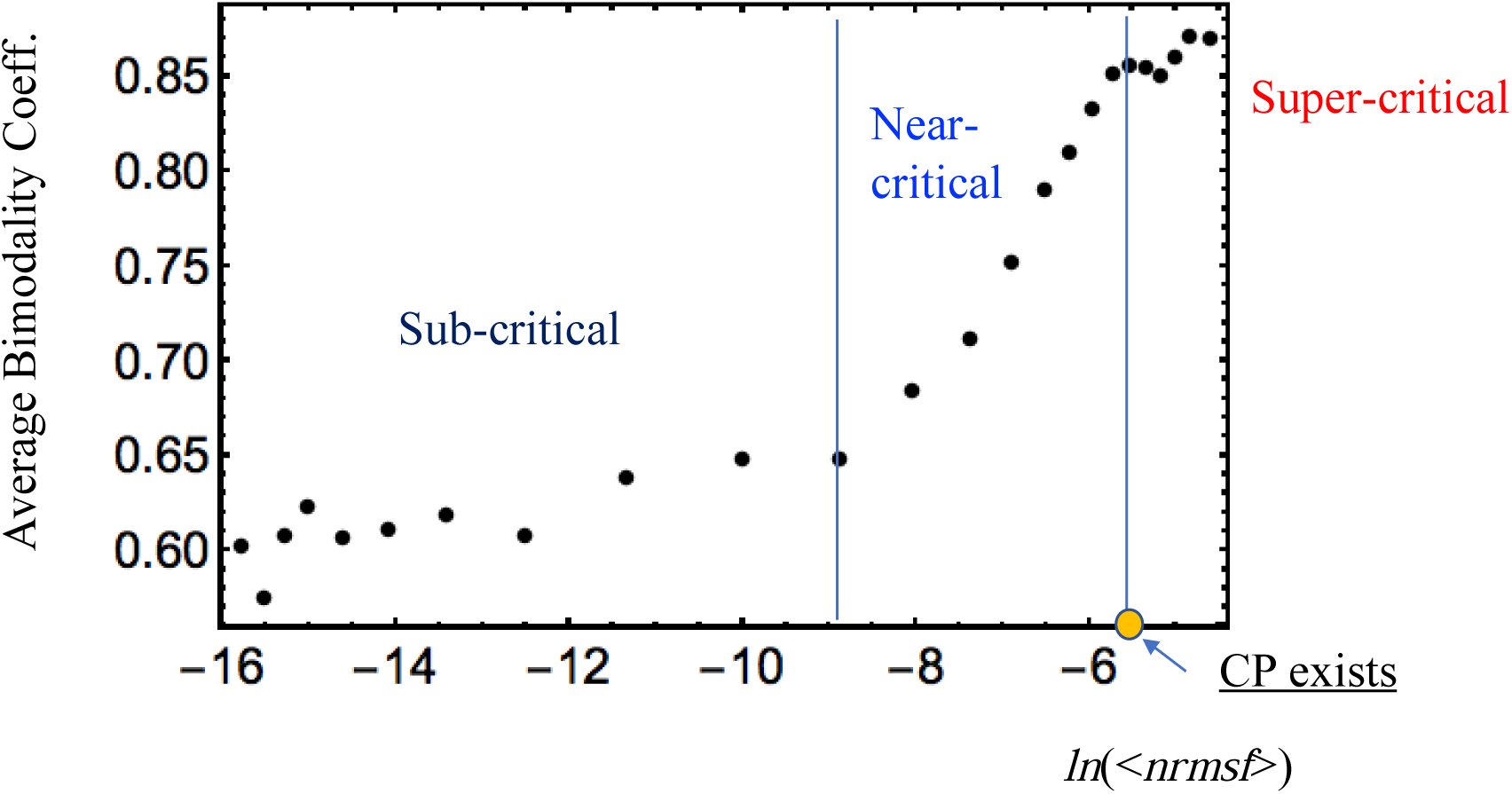
Critical states emerge through a critical transition in bimodality: Bimodality coefficients (average over all cell states) with *nrmsf* are estimated. Overall RNA expression is sorted and grouped according to the degree of *nrmsf*. This *nrmsf* grouping is made at a given sequence of discrete values (*x*_*i*_: integer and integer + 0.5 from -15 to -4) of *nrmsf* with a fixed range (*x*_*i*_ -0.5 < *x*_*i*_ < *x*_*i*_ + 0.5), and the corresponding average of the bimodality coefficient over the number of cell states is evaluated. The result clearly exhibits a transitional behavior to distinguish (averaged) critical states: with respect to the value of *ln*<*nrmsf*>, sub-critical state< -8.8; -8.8 <near-critical state < -5.4; -5.4< super-critical state, where < > indicates the ensemble average of the *nrmsf* grouping. A critical point (CP: summit of sandpile criticality: **Figure 3**) exists around the edge (*ln*(<*nrmsf*>) ∼ -5.5) between the near‐ and super-critical states.

## Results

Early embryo development involves a crucial step in global gene expression control: erasure of the initial maternal-only imprinting' of gene regulation to start a new global control that is driven by the new genetic set-up. In the zygote, the new diploid genome is still inactive in a chromatin fully condensed state, and the metabolism is orchestrated by maternal RNA molecules. Thus, for further development, it is important to erase the previous control to allow for a new start based on the new maternal/paternal mixed genome. In some sense, this ‘erasure’ path can be considered to be a transformation that proceeds in the direction opposite that for acquisition of a terminal cell fate.

**Figure 2:**
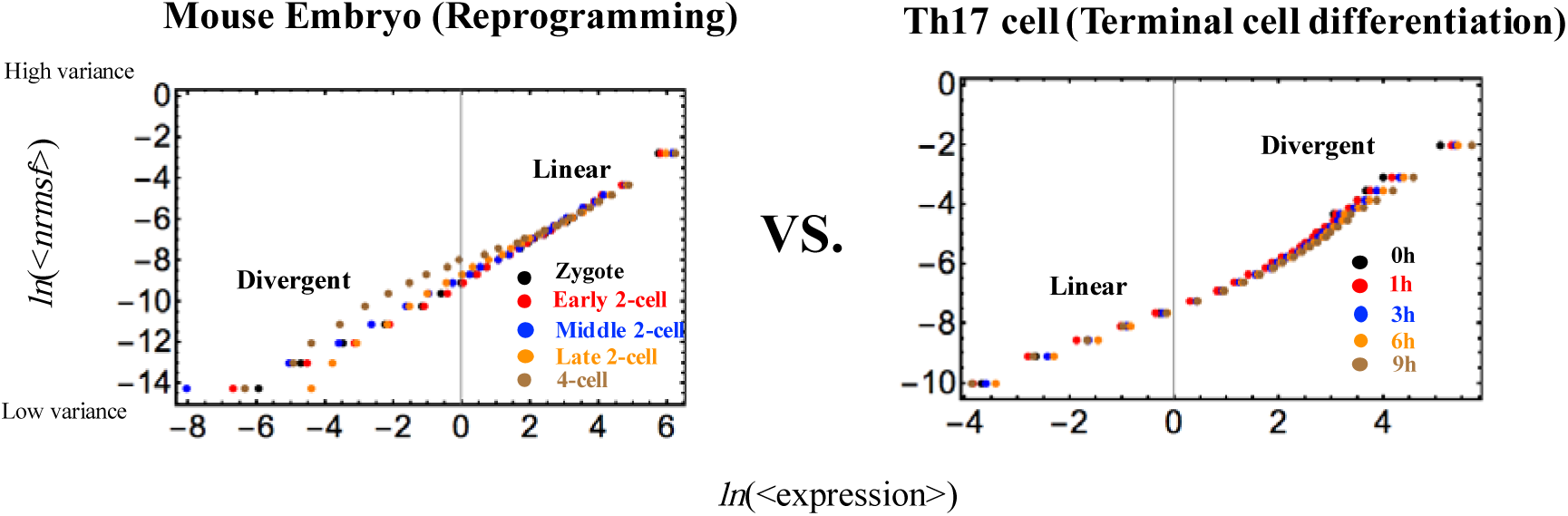
Genome Avalanches ‐ Scaling Divergent Behaviors: In overall expression, scaling-divergent behavior (genome avalanches), an important characteristic of SOC control [2,3] is seen in a log-log plot of averaging behaviors: (left panel: mouse embryo development; right panel; Th17 terminal cell differentiation). Log-log plots show the natural logarithm of the group average of expression (x-axis) and *nrmsf* (y-axis) (each dot for mouse: *n* = 685 RNAs; Th17: *n* = 525), where overall expression is sorted and grouped according to the degree of *nrmsf*. This reveals that order (scaling) and disorder (divergence) are balanced at around the CP (**Figure 3**). Two distinct biological processes (reprogramming in early mouse embryo development versus Th17 immune cell differentiation) show opposite scaling-divergent behaviors. Scaling behavior occurs in the ensemble of high-variance RNA expression (high region of *nrmsf:* supercritical state) in mouse early embryo development and divergent behavior occurs in the ensemble of low-variance RNA expression (low region of *nrmsf:* sub-critical state), whereas the Th17 terminal cell fate (single cell) shows opposite behaviors.

We found that the erasure of purely maternal expression control imprinting in the embryo can be explained by SOC and, much more importantly, the direction of early embryo development traverses the same phases of differentiation, but in the opposite order (**Figure 2**).

The erasure of criticality in global gene expression levels is a clear signature of an approaching critical transition state (**Figure 3**), providing a quantitative (open-) thermodynamic appreciation of the still largely qualitative notion of the epigenetic landscape in the reprogramming process. In mouse embryo development, the critical point (CP: a critical point at the divergence of up‐ and down-regulation) exists at the boundary of the super-critical state, whereas in differentiation, the CP is at the boundary of the sub-critical state. This points to another kind of ‘reversion’ with respect to the differentiation phase in addition to the location of the CP state.

The existence of a critical transition state is further supported by the fact that each motion in the phase space of genetic expression can be interpreted (in a completely model-free manner) as a decrease in the correlation with the initial genome-wide-expression profile with increasing time. **Figure 4** reveals such displacement dynamics for embryo early development in both human and mouse. Note that the two trajectories proceed in parallel, with only a constant delay in time for the human embryo.

**Figure 3:**
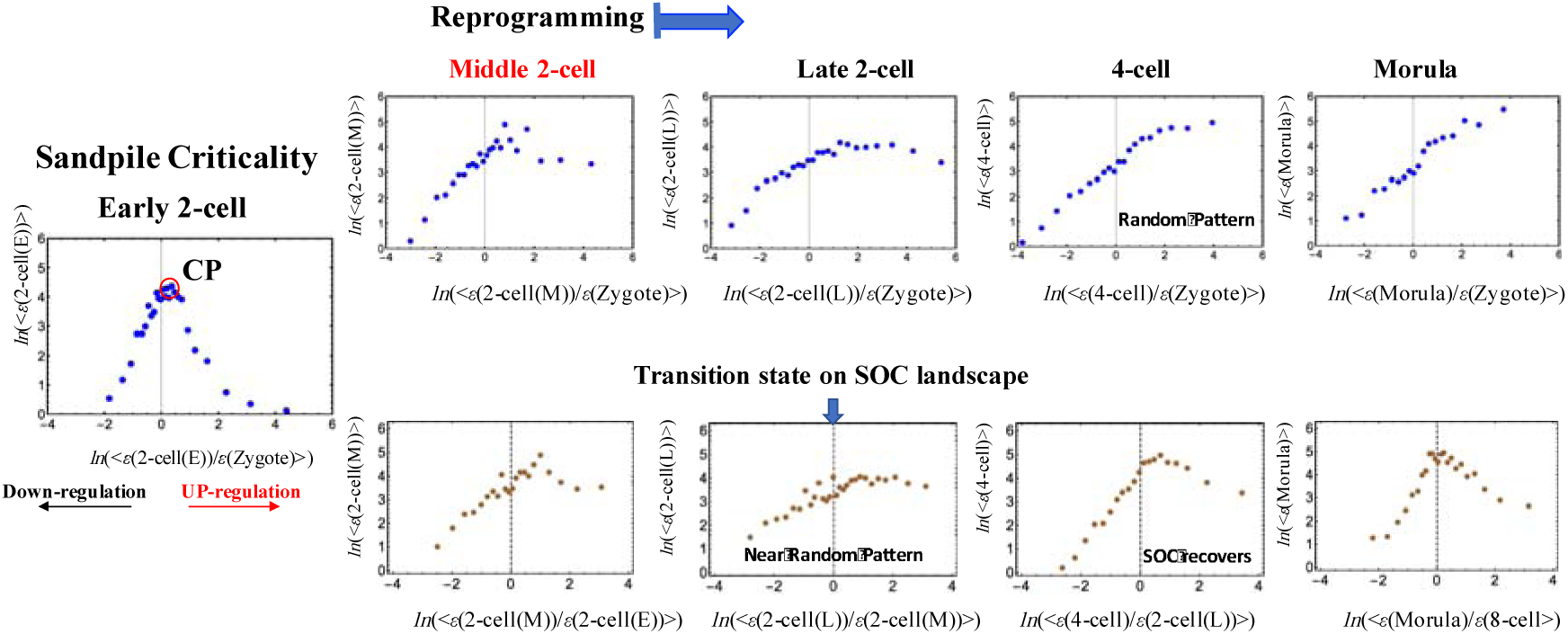
Timing of the genome-state change on the SOC control landscape in a single cell: A change in critical dynamics through sandpile-type criticality (diverging up‐ and down-regulation at the CP around *ln*(<*nrmsf*>) ∼ -5.5), which affects the entire genome expression dynamics, appears in the change in overall expression (e.g., fold-change) between different time points [2,3]. Thus, the erasure of sandpile-type criticality of the zygote state points to the timing of a genome-state change in mouse embryo development. This erasure of the zygote criticality (upper row) occurs after the middle 2-cell state to reveal a stochastic expression pattern as a linear correlative behavior (refer to [3]). The transition point occurs at the middle-later 2-cell states (lower row), at which sandpile-type criticality disappears, and thereafter recovers. This shows the existence of a SOC control landscape and a transition state at the middle-later 2-cell states. The *x*‐ and *y*-axes represent the fold-change in expression and the group average expression.

From a functional perspective, early embryo development is the ‘opposite’ of terminal cell-fate differentiation. The erasure of the previously organized state of the zygote implies an increase in stochasticity of gene expression regulation in terms of the distribution of different RNA species. This is consistent with the fact that, in the early stages of development, the embryo must eliminate the maternal-only imprinting’ of the zygote to ‘start-from-scratch’ with a new developmental program driven by the maternal/paternal genome pattern [9]. In contrast, the path toward a terminal cell fate is a path toward increasing order with respect to the undifferentiated state [10].

These findings are purely phenomenological at the present and await the discovery of mechanistic microscopic bases. However, our results suggest that a single-cell statistical thermodynamics approach to biological regulation provides useful insight to unveil the underlying mechanism on self-regulation of life in the cases where the extreme complexity of the system rules out any strict mechanistic approach.

**Figure 4:**
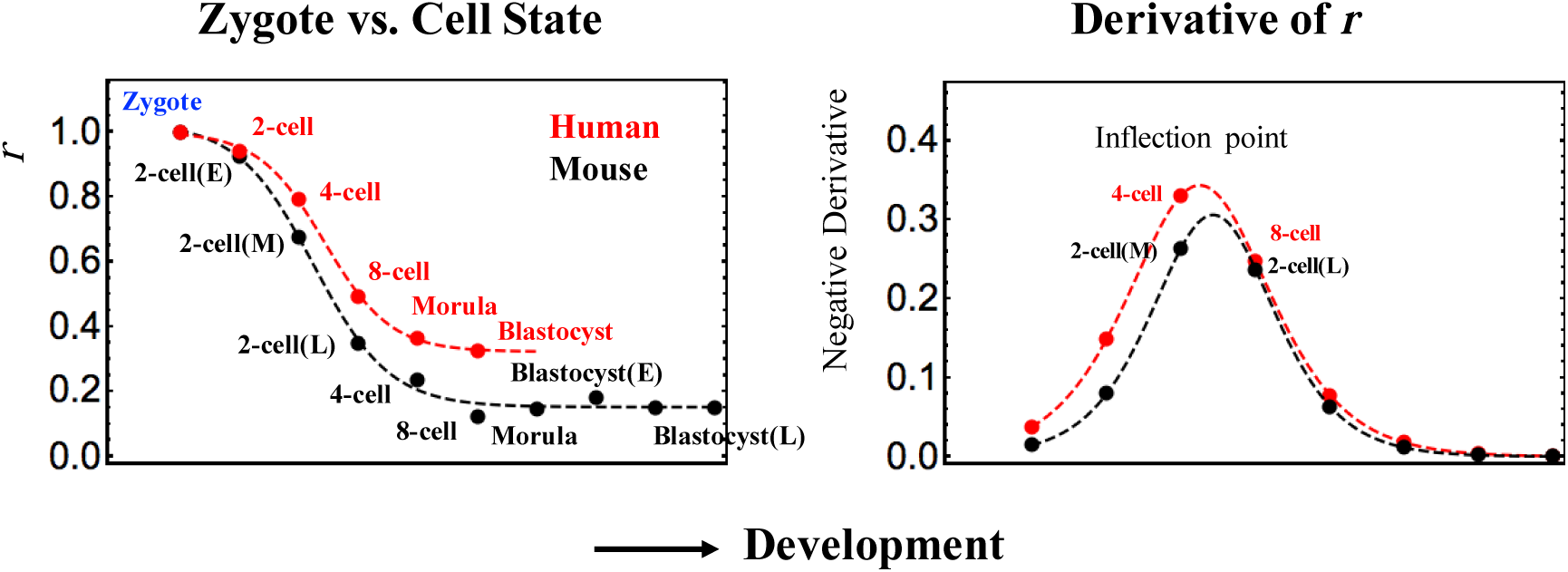
Critical transition revealed through changes in the Pearson correlation between the zygote and cell developed states: The Pearson correlation transition further supports the timing of a genome-state change via a transition state (**Figure 3**): the Pearson correlation for the cell state with the zygote exhibits a critical transition as a tangent hyperbolic function: *a* – *b* · tanh(*c* · *x* – *d*); *x* represents the cell state with *a* =0.59, *b* = 0.44 *c* = 0.78 and *d* = 2.5 (*p*< 10^−3^) for human (red dashed line), and *a* = 0.66, *b* = 0.34 *c* = 0.90 and *d* = 3.1 (*p*< 10^−2^) for mouse (black). This plot clearly shows that a phase transition occurs at the inflection point (zero second derivative of the tangent hyperbolic function), where there is a phase difference between the 4-cell and 8-cell states for human, and between the middle and late 2-cell states for mouse.

## Discussion

Building upon previous evidences of self-organized criticality (SOC) in cell fate change [1-3], we were able to get a biologically reliable description of early embryo development which provides a quantitative (open-) thermodynamic appreciation of the epigenetic landscape.

The event of (forward) reprogramming initiation corresponds to the erasure of sandpile-type critical point (CP) stemming from the initial stage of embryogenesis and gene expression around the sandpile-type CP of the initial state plays an essential role in development. The quest for the mechanistic basis of such transition is of utmost importance.

Regarding backward single cell reprogramming such as an induced pluripotent stem (iPS) cell from a somatic cell, a stochastic model [11,12] has shown how the success of the reprogramming depends upon the ectopic expression of transcription factors such as Yamanaka factors in a probabilistic manner. The stochastic model can be due to temporal-spatial asynchronous molecular reaction events in heterogeneous cell populations [13]. Thus, investigation of the breakdown of SOC control in reprogramming of single adult somatic cell will be important for understanding how and why a single cell can overcome an epigenetic landscape potential barrier to succeed the backward reprogramming. A comparative analysis in the breakdown of SOC control between success and failure of reprogramming in single adult cell could elucidate underlying molecular mechanism to explain why single somatic cell succeeds or fails reprogramming in the genome. Here it is expected that (no) breakdown of SOC control occurs in the (failure) success of single cell reprogramming or multiple steps of breakdown of SOC control might exist. Hence, findings will be fundamentally different from current trial-and-error combinatorial approaches in terms of the combination of activating enhancers and inhibiting barriers of reprogramming [14]. Furthermore, this might lead to understanding of distinct novel molecular mechanism in change from single cell to population of cells.

## Methods

### Biological Data Sets

We analyzed mammalian RNA-Seq data:

i)Early embryonic development in human and mouse developmental stages in RPKM (Reads Per Kilobase Mapped) values; GEO ID: GSE36552 (human: *N* = 20286 RNAs) and GEO ID: GSE45719 (mouse: *N* = 22957 RNAs), which have 7 and 10 embryonic developmental stages (experimental details in [15] and [16], respectively):

Human: oocyte (m=3), zygote (m=3), 2-cell (m=6), 4-cell (m=12), 8-cell (m=20), morula (m=16) and blastocyst (m=30), Mouse: zygote (*m*=4), early 2-cell (*m*=8), middle 2-cell (*m*=12), late 2-cell (*m*=10), 4-cell (*m*=14), 8-cell (*m*=28), morula (*m*=50), early blastocyst (*m*=43), middle blastocyst (*m*=60) and late blastocyst (*m*=30), where *m* is the total number of single cells.

ii) T helper 17 cell differentiation from mouse naive CD4+ T cells in RPKM values, where Th17 cells are cultured with anti-IL4, anti-IFN_γ_, IL-6 and TGF-β (details in [17]); GEO ID: GSE40918 (mouse: *N* = 22281 RNAs), which has 9 time points: *t*_0_ = 0, *t*_1_ = 1,3,6,9,12,16,24, *t*_*T*=8_ = 48h.

RNAs that had RPKM values of 0 over all of the cell states were excluded. Random real numbers in the interval [0-1] generated from a uniform distribution were added to all expression values for the natural logarithm. This procedure avoids the divergence of zero values in the logarithm. The robust sandpile-type criticality through the grouping of expression (**Figure 1B**) was checked by multiplying the random number by a positive constant, *a* (*a*< 10), and we set *a* = 0.01. Note: The addition of large random noise (a>>10) destroys the sandpile CP.

### SOC Control Mechanism of the Cell-fate Change

We previously demonstrated that the critical dynamics of self-organizing whole-genome expression (self-organized criticality: SOC) play essential roles in determining the cell-fate change in distinct biological processes at both the single-cell and population (tissue) levels [3].

In SOC-controlled self-organization (in terminal cell differentiation), *nrmsf* (normalized root mean square fluctuation) acts as an order parameter for the self-organization of whole gene expression [2,3]. We can examine if *nrmsf* acts as an order parameter in early embryo development, which is determined by dividing *rmsf* by the maximum overall {*rmsf*_*i*_}:

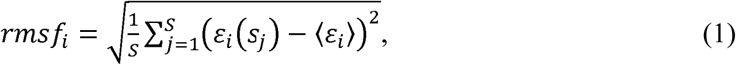
where *rmsf*_*i*_ is the *rmsf* value of the *i*^*th*^ RNA expression, which is expressed as *ε*_*i*_(*s*_*j*_) at a specific cell state *s*_*j*_ (e.g., in mouse, *S* = 10: *s*_1_ = zygote, early 2-cell, middle 2-cell, late 2-cell, 4-cell, 8-cell, morula, early blastocyst, middle blastocyst and *s*_10_ = late blastocyst), and 〈*ε*_*i*_〉 is its expression average over the number of cell states. Regarding self-organization with critical dynamics, we investigated averaging behaviors in *nrmsf*, and in the fold-change in expression, where we observed a stochastic resonance effect in the terminal cell fate process. For methodological details, refer to our previous works [2,3].

## Acknowledgments

MT sincerely thanks the Institute for Advanced Biosciences, Keio University, Tsuruoka City, the Yamagata prefectural government, Japan, and Mr. Fumiaki Kikuchi for allowing him to complete this research project at Keio University. The authors are thankful to Dr. Midori Hashimoto for helping to produce the Figures.

